# Cannabidiol Reduces Oxycodone Self-Administration While Preserving Its Analgesic Efficacy in a Rat Model of Neuropathic Pain

**DOI:** 10.1101/2025.04.21.649778

**Authors:** Adriaan W. Bruijnzeel, Azin Behnood-Rod, Ranjithkumar Chellian, Wendi Malphurs, Robert M. Caudle, Marcelo Febo, Barry Setlow, Niall P. Murphy, John K. Neubert

**Affiliations:** Department of Psychiatry, University of Florida, Gainesville, FL, USA; Center for Addiction Research and Education, University of Florida, Gainesville, USA; Department of Orthodontics, University of Florida, Gainesville, FL, USA; Department of Oral and Maxillofacial Surgery, University of Florida, Gainesville, FL, USA; Department of Biomedical Sciences, Texas A&M University, Dallas, TX, USA

**Author notes:** Corresponding author: Adriaan Bruijnzeel, PhD, University of Florida, Department of Psychiatry, 1149 Newell Dr., Gainesville, Florida 32611.

## Abstract

Prescription opioid misuse is a significant public health concern among individuals with chronic pain. Treating severe pain often requires high doses of opioids, increasing the risk of developing an opioid use disorder. Cannabidiol (CBD) is a non-psychoactive component of cannabis that has shown therapeutic potential without abuse liability. This study investigated the effects of CBD on oxycodone self-administration and hyperalgesia in an animal model of chronic neuropathic pain. Adult male rats were trained to self-administer intravenous oxycodone (0.06 mg/kg/infusion). Subsequently, they underwent chronic constriction injury (CCI) of the sciatic nerve or received sham surgery. Paw withdrawal latency was measured using the Hargreaves test as an indicator of thermal pain sensitivity. CBD (0, 1, 3, and 10 mg/kg, IP) was administered before the self-administration sessions, and pain testing was conducted afterward. The rats acquired oxycodone self-administration, as indicated by more active than inactive lever presses. CCI surgery decreased the paw withdrawal latency, confirming the induction of neuropathic pain. CCI alone did not affect oxycodone self-administration, suggesting that neuropathic pain does not affect opioid intake. Treatment with CBD reduced oxycodone self-administration in both the sham and CCI rats. Oxycodone self-administration in the CCI rats reversed the CCI-induced decrease in paw withdrawal latency. However, CBD did not affect the antinociceptive effect of oxycodone in CCI rats. Taken together, these findings demonstrate that CBD reduces oxycodone self-administration without affecting the antinociceptive effects of oxycodone in neuropathic pain. This study supports the potential of CBD to reduce opioid use and misuse, regardless of pain status.

## 1. Introduction

The use of prescription opioids is widespread in the United States. It has been estimated that between 2019 and 2020, twelve percent of the general population used prescription opioids (Zajacova et al., 2023). This prevalence increases among individuals with pain conditions. Thirty percent of those with chronic pain and more than forty percent of those with high-impact chronic pain report using prescription opioids (Zajacova et al., 2023), highlighting that opioids continue to play a critical role in pain management. It has been estimated that twelve percent of the adults with an opioid prescription misuse opioids (Han et al., 2024). The motives for opioid misuse vary by age. Younger individuals often misuse opioids to experiment, relax, get high, or cope with negative emotions. In contrast, older adults typically misuse opioids for pain relief (Schepis et al., 2019). During the last two decades, there has been a strong increase in drug overdose deaths. The number of overdose deaths increased from fewer than 20,000 in 1999 to more than 100,000 in 2023 (Centers for Disease Control and Prevention, 2021; 2024). The great majority of these deaths are caused by synthetic opioids (Seth et al., 2018; Spencer et al., 2024). Therefore, it is critical to develop treatment approaches with potential to reduce opioid use and misuse.

Cannabis is widely used for the treatment of pain, but it also has psychoactive effects that produce abuse liability (Zehra et al., 2018). Cannabis contains a complex profile of cannabinoids, which have different pharmacological actions and therapeutic potential. Tetrahydrocannabinol (THC) is the main psychoactive constituent that produces the euphoric effects of cannabis and is responsible for its abuse liability (Ashton, 2001). Conversely, cannabidiol (CBD) exhibits a wide range of therapeutic properties, including anti-inflammatory, anxiolytic, and anticonvulsant effects, but is generally considered devoid of psychoactive effects (Mechoulam et al., 2002; Izzo et al., 2009; Starowicz and Di Marzo, 2013).

Cannabinoids exert their effects primarily by binding to the CB1 receptor, which is highly expressed in the brain, but also present at lower levels in peripheral organ systems (Haspula and Clark, 2020). They also bind to the CB2 receptor, which is highly expressed in immune cells and in lymphoid tissues (Haspula and Clark, 2020). CBD is a negative allosteric modulator of the CB1 receptor and an orthosteric partial agonist at the CB2 receptor (Laprairie et al., 2015; Tham et al., 2019). CBD has also been shown to act as a positive allosteric modulator at GABA-A receptors (Bakas et al., 2017a). CBD is further believed to enhance levels of endocannabinoids such as anandamide by inhibiting the enzyme fatty acid amide hydrolase (FAAH), which breaks down anandamide (Peng et al., 2022). Increased anandamide levels can lead to reduced pain perception (Clapper et al., 2010). Cannabidiol has exhibited analgesic properties in various pre-clinical animal models of pain as well as in clinical trials (Silva-Cardoso and Leite-Panissi, 2023)

Chronic use of opioids can lead to tolerance, physical dependence, and withdrawal symptoms upon discontinuation (Bruijnzeel et al., 2006; Liu et al., 2008; Volkow and McLellan, 2016). High-potency opioids like oxycodone are often used for their effective pain relief, but they also have a significant risk of abuse (Remillard et al., 2019; Kibaly et al., 2021). Oxycodone is a semi-synthetic opioid analgesic that is chemically derived from the naturally occurring opioid thebaine. Oxycodone activates mu, delta, and kappa 2b-opioid receptors and has a higher affinity for mu-opioid receptors (Ki value, 18 nM) than for delta (Ki value, 958 nm) and kappa-opioid receptors (Ki value, 677 nm)(Monory et al., 1999; Nielsen et al., 2007; Yang et al., 2016). Oxycodone’s analgesic effects result from activating mu-opioid receptors in the brain, but stimulation of these receptors also causes euphoria, anxiolysis, and sedation (Stein, 2013; Valentino and Volkow, 2018; Bruijnzeel et al., 2022). Due to its potent efficacy for pain relief, oxycodone is widely used in clinical settings, but chronic use leads to tolerance and dependence. Given the high potential for misuse associated with oxycodone use for pain treatment, exploring adjunct therapies that could mitigate this risk is essential. Therefore, in the present study, we investigated the effects of CBD on oxycodone self-administration and its effect on thermal pain sensitivity in an animal model of chronic neuropathic pain.

## 2. Materials and methods

### 2.1. Animals

Adult male Sprague Dawley rats (SD rats; 280-350 g; 8-9 weeks of age; N=24) were purchased from Charles River (Raleigh, NC). The rats were kept in standard housing conditions (2 rats per cage) in a climate-controlled vivarium on a 12 h light-dark cycle (light off at 7 PM). The study was conducted during the light period of the light-dark cycle between 10 AM and 4 PM. Food and water were available ad libitum in the home cage. The experimental protocols were approved by the University of Florida Institutional Animal Care and Use Committee.

### 2.2. Drugs and treatment

For intravenous self-administration, oxycodone hydrochloride (NIDA Drug Supply Program) was dissolved in sterile saline (0.9 % sodium chloride). The rats self-administered 0.06 mg/kg/infusion of oxycodone in a volume of 0.1 ml/infusion. The oxycodone dose is expressed as the weight of the salt. CBD (NIDA Drug Supply Program) was prepared in 5 % ethanol and 5 % Cremophor in PBS and administered intraperitoneally (IP) at a volume of 1 ml/kg body weight.

### 2.3. Experimental design

We investigated the effect of CBD on oxycodone self-administration in male rats under conditions of chronic pain induced by chronic constriction injury (CCI) and in non-pain (sham) states. A schematic overview of the experimental design is depicted in Figure 1. Twenty-four rats were trained to press a lever for food pellets over a 10-day period. One week after the food training sessions, all rats underwent jugular vein catheterization surgery over 4 days to enable intravenous self-administration (IVSA) of oxycodone. Following surgery, the animals were given at least 7 days to recover. After recovery, a food reminder session was conducted using a fixed-ratio 1 (FR1) schedule with a 20-second time-out (TO20) during a 20-minute session to confirm that the rats retained lever-pressing behavior. The rats were then trained to acquire oxycodone self-administration (0.06 mg/kg/infusion) on an FR1-TO20 schedule during 120-minute sessions for 14 sessions. The left lever served as the active lever for 11 rats, while the right lever served as the active lever for 12 rats. The active lever assignment remained the same throughout the study. Responding on the active lever resulted in the delivery of an oxycodone infusion (0.1 ml infused over a 6.5-s period). The infusion was paired with a cue light, which remained illuminated throughout the time-out period. Responding on the inactive lever was recorded but did not have scheduled consequences. The active and inactive lever were retracted during the time-out period. Self-administration sessions were conducted five days per week. Following acquisition of oxycodone self-administration, rats were divided into two surgical groups: sham (N=11) and CCI (N=11). CCI surgery was performed to induce neuropathic pain, while sham surgery was performed to serve as a control. The CCI and sham groups were counterbalanced based on the left/right side of the active lever. In the CCI group, six rats self-administered oxycodone with the right lever as the active lever, while five rats used the left lever as the active lever. In the sham group, five rats self-administered oxycodone with right lever as the active lever, while six rats used the left lever as the active lever. Previous studies have shown that CCI of the sciatic nerve induces thermal hypersensitivity in adult male SD rats, as measured by the Hargreaves test, which remains relatively stable for at least one month post-surgery (Baliki et al., 2005; Guillemette et al., 2012; Jørgensen et al., 2012; Xi et al., 2023). In this study, the Hargreaves test, oxycodone self-administration, and CBD treatment were conducted between post-CCI days 5 and 20. After the post-operative recovery period of four days, the Hargreaves test was conducted to assess thermal nociception as a measure of pain sensitivity in sham and CCI rats on post-CCI day 5. Baseline oxycodone self-administration sessions were then conducted for 5 days, with the Hargreaves test performed in the morning prior to the day 4 and 5 baseline self-administration sessions (i.e. on post-CCI day 10 and 11). Following the five-baseline oxycodone self-administration sessions, post-CCI day 12 – 20, CBD (0, 1, 3, and 10 mg/kg, IP) was administered according to a Latin square design, with 48 h separating successive CBD sessions. Twenty minutes after the CBD injection, the rats underwent a 2-hour oxycodone self-administration session. Immediately following the self-administration session, the Hargreaves test was performed to assess any changes in pain sensitivity due to CBD treatment. Self-administration was also assessed in the intervening sessions (between successive CBD sessions). This experimental design allowed us to evaluate the effects of different doses of CBD on oxycodone intake in sham control rats and rats with chronic pain, as well as to assess any potential interaction effects of CBD and oxycodone on thermal hyperalgesia induced by CCI. One rat was excluded from the study after implantation of catheter due to health issues. Additionally, one rat from the CCI group was excluded from the study due to excessively high responding on the inactive lever (> 200 inactive lever responses) across post-CCI self-administration sessions. During the self-administration period, catheter patency was tested by infusing 0.2 ml of the ultra-short-acting barbiturate Brevital (1% methohexital sodium). Rats with patent catheters displayed a sudden loss of muscle tone. If the rats did not respond to Brevital, their self-administration data were excluded from the analysis. On day 10 of oxycodone self-administration acquisition, one rat did not respond to Brevital, and its data were excluded.

**Figure 1.**
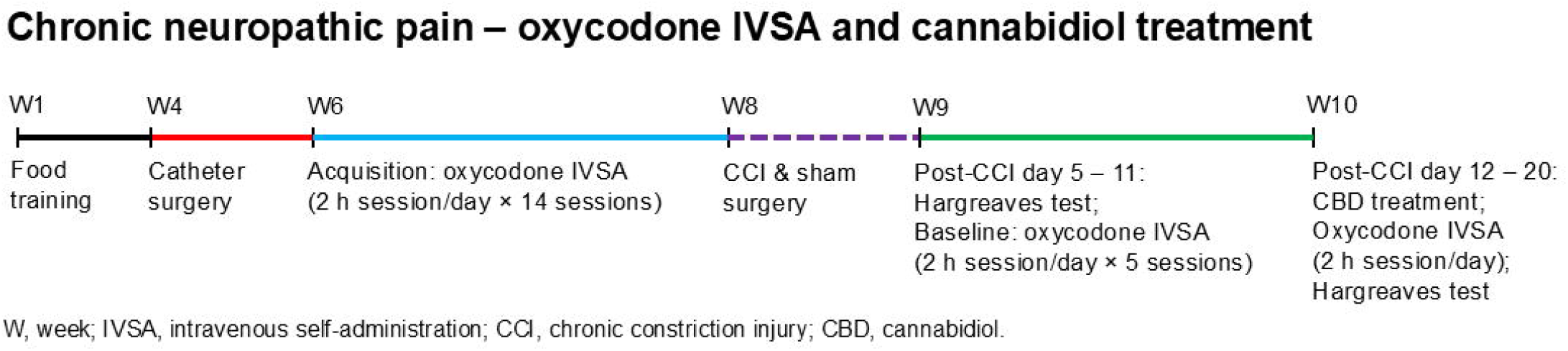
Schematic overview of the experiment. Adult male Sprague Dawley rats were trained to respond for food pellets. Following this, the rats were implanted with IV catheters and trained to self-administer oxycodone in 2-hour sessions per day for 14 sessions. After the acquisition phase, chronic constriction injury (CCI) surgery was performed to induce neuropathic pain, while sham surgery was performed in control animals. Following recovery, the Hargreaves test was used to assess thermal pain sensitivity, and baseline oxycodone self-administration sessions were conducted in 2-hour sessions per day for 5 sessions. Subsequently, the effects of cannabidiol (CBD) treatment on oxycodone self-administration and thermal pain sensitivity were evaluated in both CCI and sham control rats. Abbreviation: IVSA, intravenous self-administration.

### 2.4. Food training

Rats were trained to press a lever for food pellets in operant chambers placed in sound- and light-attenuated cubicles (Med Associates, St. Albans, VT) as in our previous work (Zislis et al., 2007; Yamada and Bruijnzeel, 2011). Food delivery was paired with a cue light, which remained illuminated throughout the time-out period. Twenty-four male rats were initially trained to respond for food pellets (F0299, 45 mg, chocolate-flavored pellets; Bio-Serv, Flemington, NJ) on a FR1 schedule using both levers over a period of 5 days. This was followed by an additional 5 days of training on an FR1 schedule with a 20-second time-out during 20-minute sessions using both levers. After completing the food training session on day 8, the rats were singly housed and remained singly housed for the remainder of the study. Three days before the start of the oxycodone self-administration sessions, the rats were allowed to respond for food pellets under the FR1-TO20 schedule during a single 20-minute session. Responding on both the right and left levers resulted in the delivery of a food pellet.

### 2.5. Intravenous catheter implantation

The catheters were implanted as described before (Chellian et al., 2021a; Chellian et al., 2021b; Chellian et al., 2022). The rats were anesthetized with an isoflurane-oxygen vapor mixture (1-3%) and prepared with a catheter in the right jugular vein. The catheters consisted of polyurethane tubing (length 10 cm, inner diameter 0.64 mm, outer diameter 1.0 mm, model 3Fr, Instech Laboratories, Plymouth Meeting, PA). The right jugular vein was isolated, and the catheter was inserted 2.9 cm. The tubing was then tunneled subcutaneously (SC) and connected to a vascular access button (Instech Laboratories, Plymouth Meeting, PA). The button was exteriorized through a 1-cm incision between the scapulae. During the 7-day recovery period, the rats received daily infusions of the antibiotic Gentamycin (4 mg/kg, IV, Sigma-Aldrich, St. Louis, MO). A sterile heparin solution (0.1 ml, 50 U/ml) was flushed through the catheter before and after administering the antibiotic and after oxycodone self-administration. After flushing the catheter, 0.05 ml of a sterile heparin/glycerol lock solution (500 U/ml, Instech Laboratories, Plymouth Meeting, PA) was infused into the catheter. The animals received carprofen (5 mg/kg, SC) daily for 72 hours after the surgery.

### 2.6. CCI surgery

Chromic gut suture (4.0, Ethicon) was cut into 2 cm pieces and immersed in sterile saline to prevent drying. Surgery was performed under aseptic conditions on a heating pad. Animals were administered buprenorphine (1.0 mg/kg, SC) before being maintained under general anesthesia (1-3% isoflurane in oxygen). The right hind leg was shaved and sterilized with chlorohexidine and 70% ethanol. A 5 to 7 mm incision was made in the skin below the femur and the skin separated from the muscle and connective tissue using blunt forceps. An incision was made through femoris muscles, and the sciatic nerve freed using a glass pipette with a blunt curved tip. Three ligatures were made using a double knot 1 mm apart, tightened until the loop was just snug and the ligatures were unable to slide along the nerve. Silk sutures (5.0, Ethicon) were used to close the muscle layer and skin, and the wound was cleaned with chlorohexidine before application of triple antibiotic ointment. Rats in the sham control group underwent the same procedure (i.e., same paw, surgical intervention, treatment etc.) with the exception that no ligatures were placed.

### 2.7. Hargreaves thermal paw withdrawal test

Animals were acclimated to a thermal hindpaw reflex testing apparatus (Ugo Basile model 7375a, Stoelting, Wood Dale, IL, USA) until they exhibited minimal spontaneous movement. A radiant heat lamp (intensity set at 45) was aimed at the left or right hind paw (random order) until the paw was withdrawn, with a cutoff time of 25 seconds to prevent tissue damage. The average of two latency measurements was used for analysis.

### 2.8. Statistics

Acquisition of oxycodone self-administration (14 sessions) data were analyzed using two-way ANOVAs with lever (active versus inactive) and session as within-subject factors. Oxycodone self-administration data (5 sessions) following sham and CCI surgery were analyzed using three-way ANOVAs with lever and session as within-subject factors, and surgery (sham versus CCI) as a between-subject factor. Hargreaves test data collected post-surgery, including assessments before the day 4 and 5 baseline oxycodone self-administration sessions, were analyzed using two-way ANOVAs with paw (right versus left) as a within-subject factor and surgery as a between-subject factor. Post-surgery, CBD treatment effects on oxycodone self-administration were analyzed using two-way ANOVAs with treatment as a within-subject factor and surgery as a between-subject factor. CBD treatment effects on Hargreaves test data (absolute paw withdrawal latency and percentage change from baseline) collected after oxycodone self-administration were analyzed using three-way ANOVAs with treatment and paw as within-subject factors, and surgery as a between-subject factor. The paw withdrawal latencies were expressed as a percentage of the drug-free baseline values obtained from the Hargreaves test conducted on post-CCI surgery days 10 and 11, prior to the initiation of CBD and oxycodone self-administration. For all statistical analyses, significant interaction effects identified in the ANOVAs were followed by Bonferroni’s post hoc tests to determine which groups differed. P-values less than or equal to 0.05 were considered significant. Data were analyzed using SPSS Statistics version 29 and GraphPad Prism version 10.1.2. Figures were generated using GraphPad Prism version 10.1.2.

## 3. Results

### 3.1. Acquisition of oxycodone self-administration

The rats were initially allowed to self-administer oxycodone for 14 sessions. During this period, the rats responded more on the active lever than on the inactive lever and responding on the inactive lever decreased more over time than responding on the active lever (Fig. 2A, Lever F1,20=9.134, P < 0.01; Session F13,260=6.293, P < 0.001; Lever × Session F13,260=6.253, P < 0.001). The post hoc analysis revealed that the active lever responses were significantly higher than inactive lever responses from the eleventh session onward (Fig. 2A). In addition, inactive lever presses were significantly reduced from the second session onward compared to the first session (Fig. 2A). Furthermore, oxycodone intake initially decreased and then remained relatively stable over time (Fig. 2B, Session F13,260=4.072, P < 0.001). The post hoc analysis showed that oxycodone intake was significantly lower on the fifth session compared to the first session (Fig. 2B).

**Figure 2.**
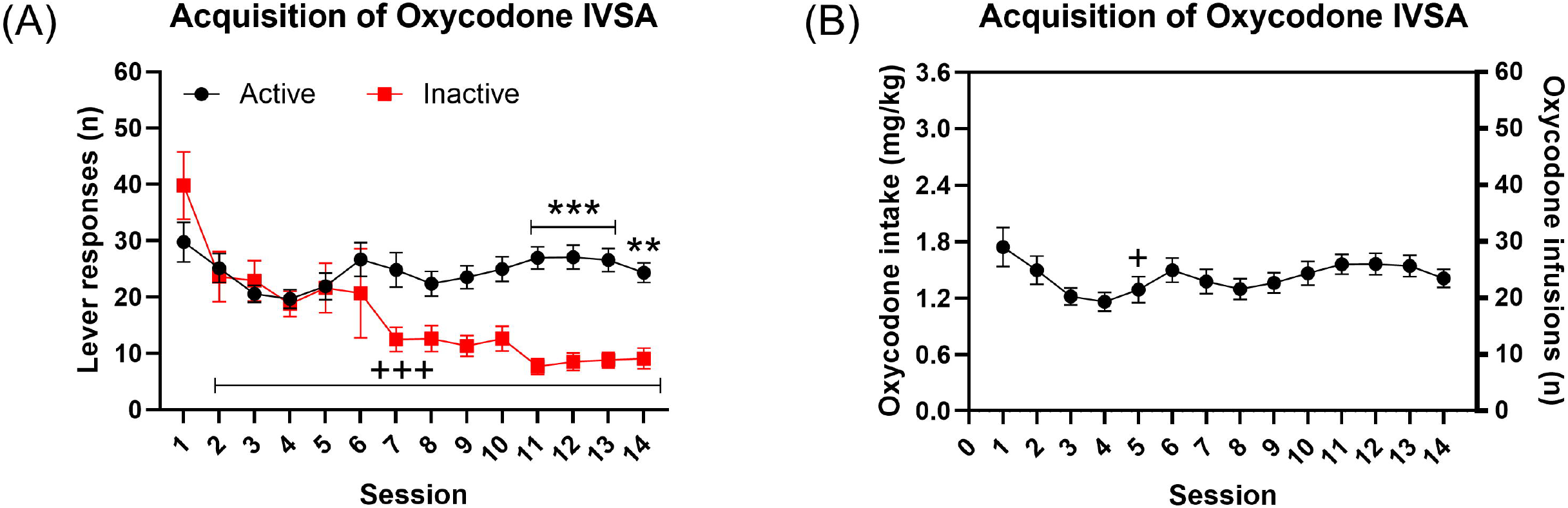
Acquisition of oxycodone self-administration in male rats. The rats were trained to respond for food pellets and then given access to oxycodone. The rats self-administered oxycodone for 14 sessions and active and inactive lever presses (A) and oxycodone intake (B) are shown. Asterisks indicate a significant difference in active lever responses and inactive lever responses during the same self-administration session. Plus sign indicate fewer left lever responses or lower oxycodone intake compared with the first self-administration session. +, P<0.05; **, P<0.01; +++, ***, P<0.001. N=21. Data are expressed as mean ± SEM.

### 3.2. CCI and Hargreaves test

The CCI surgery decreased the paw withdrawal latency, and this effect was greater in the CCI paw than in the control paw (Fig. 3A, Surgery F1,19=5.583, P < 0.05; Paw F1,19=0.514, NS; Paw × Surgery F1,19=7.126, P < 0.05). However, the post hoc analysis did not reveal a significant difference in paw withdrawal latency between the control paw and surgery paw in sham and CCI rats (Fig. 3A). Furthermore, on post-CCI days 10 and 11 (before oxycodone self-administration sessions), withdrawal latency was shorter in the CCI paw compared to the paw that did not receive surgery and both paws of animals in the sham control group (Fig. 3B, post-CCI day 10: Surgery F1,19 = 7.872, P < 0.05; Paw F1,19 = 11.926, P < 0.01; Paw × Surgery F1,19 = 11.229, P < 0.01; Fig. 3C, post-CCI day 11: Surgery F1,19 = 4.475, P < 0.05; Paw F1,19 = 16.578, P < 0.001; Paw × Surgery F1,19 = 21.592, P < 0.001). On both these days, the post hoc analysis showed that only in CCI rats the paw withdrawal latency was significantly lower in surgery paw compared with control paw (Fig. 3B and 3C).

**Figure 3.**
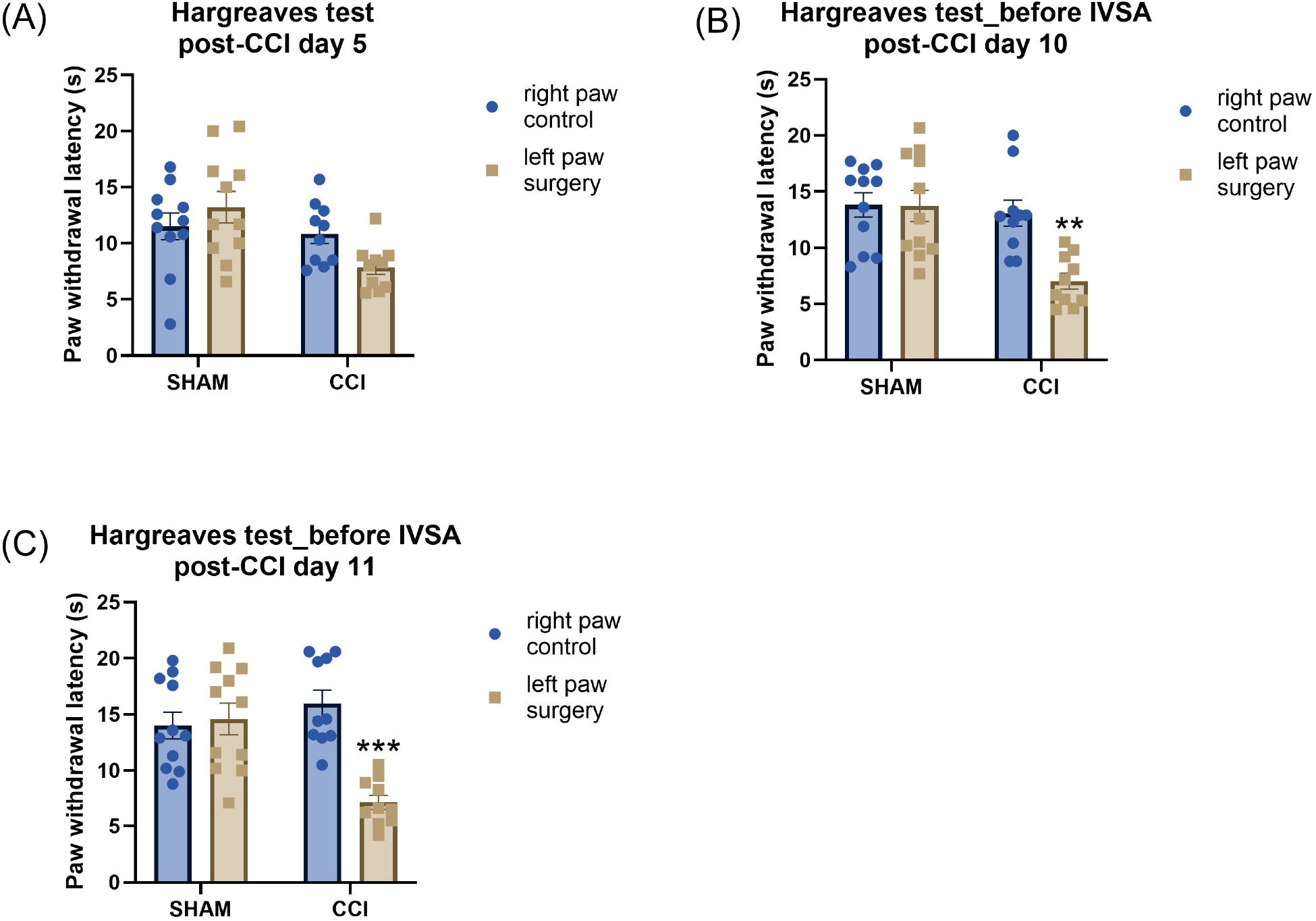
Paw withdrawal latencies are reduced in the Hargreaves test following CCI surgery. Thermal withdrawal latencies were measured in sham control and CCI rats on post-CCI day 5 (A), prior to the fourth (B, post-CCI day 10) and fifth (C, post-CCI day 11) baseline oxycodone self-administration sessions. Asterisks indicate lower paw withdrawal latencies in CCI surgery paw compared with control paw in CCI rats. **, P<0.01; ***, P<0.001. Sham, N=11; CCI N=10. Data are presented as mean ± SEM.

### 3.3. CCI and oxycodone self-administration

The CCI surgery did not affect responding on the active or the inactive lever, and the rats continued to respond more on the active than the inactive lever (Fig. 4A, Surgery F1,19=0.026, NS; Lever F1,19=43.123, P < 0.001; Lever × Surgery F1,19=0.001, NS; Session F4,76=0.466, NS; Session × Surgery F4,76=1.524, NS; Lever × Session F4,76=2.677, P < 0.05; Lever × Session × Surgery F4,76=1.228, NS). The CCI surgery also did not affect oxycodone intake (Fig. 4B, Surgery F1,19=0.010, NS; Session F4,76=1.077, NS; Session × Surgery F4,76=1.778, NS).

**Figure 4.**
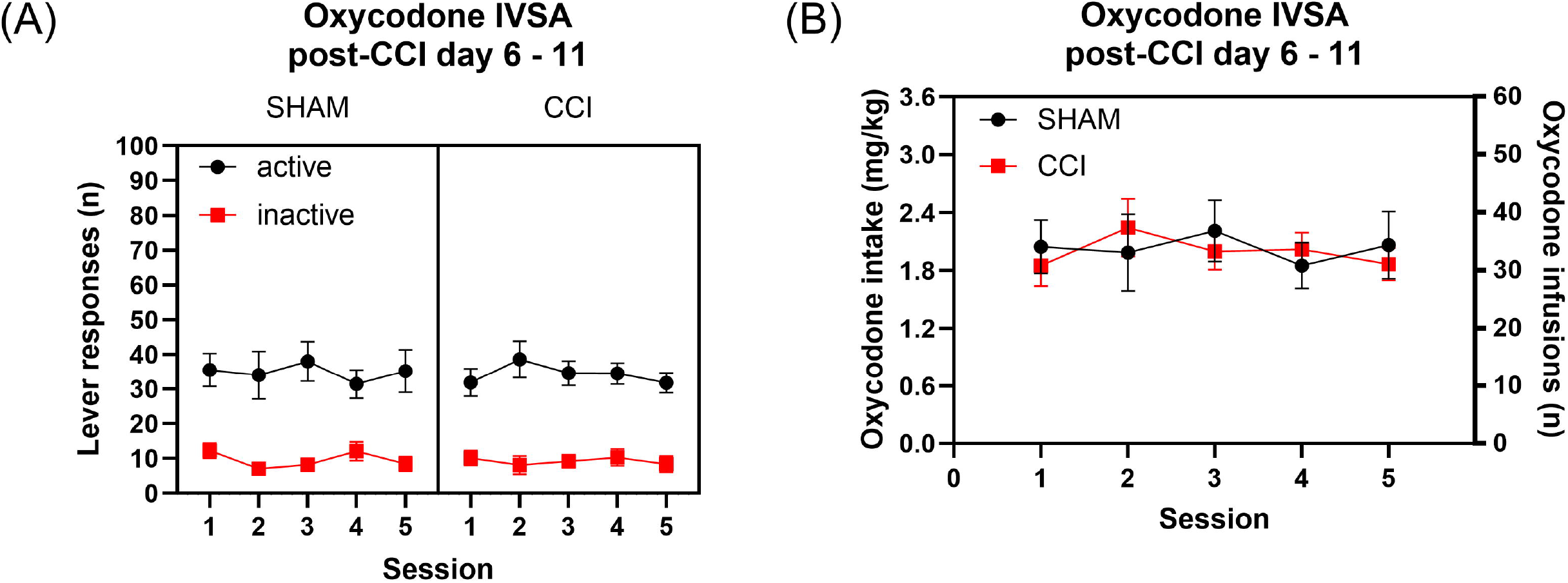
Neuropathic pain does not affect oxycodone self-administration in rats. The figures show oxycodone self-administration in sham control rats and CCI rats. The rats self-administered oxycodone for 5 days after the sham or CCI surgery and active and inactive lever presses (A) and oxycodone intake (B) are shown. Sham, N=11; CCI N=10. Data are expressed as mean ± SEM.

### 3.4. CBD treatment: 20 minutes after administration

#### 3.4.1. CBD and oxycodone self-administration

Treatment with CBD decreased responding on the active lever in both the sham control and CCI group and the magnitude of this reduction did not differ between the two groups (Fig. 5A, Surgery F1,19=1.072, NS; Treatment F3,57=5.151, P < 0.01; Treatment × Surgery F3,57=2.045, NS). Neither CBD nor surgery condition affected inactive lever responses (Fig. 5B, Surgery F1,19=0.033, NS; Treatment F3,57=0.109, NS; Treatment × Surgery F3,57=0.556, NS). CBD also decreased oxycodone intake to an equivalent extent in both surgical conditions (Fig. 4C, Surgery F1,19=1.266, NS; Treatment F3,57=4.614, P < 0.01; Treatment × Surgery F3,57=1.832, NS).

**Figure 5.**
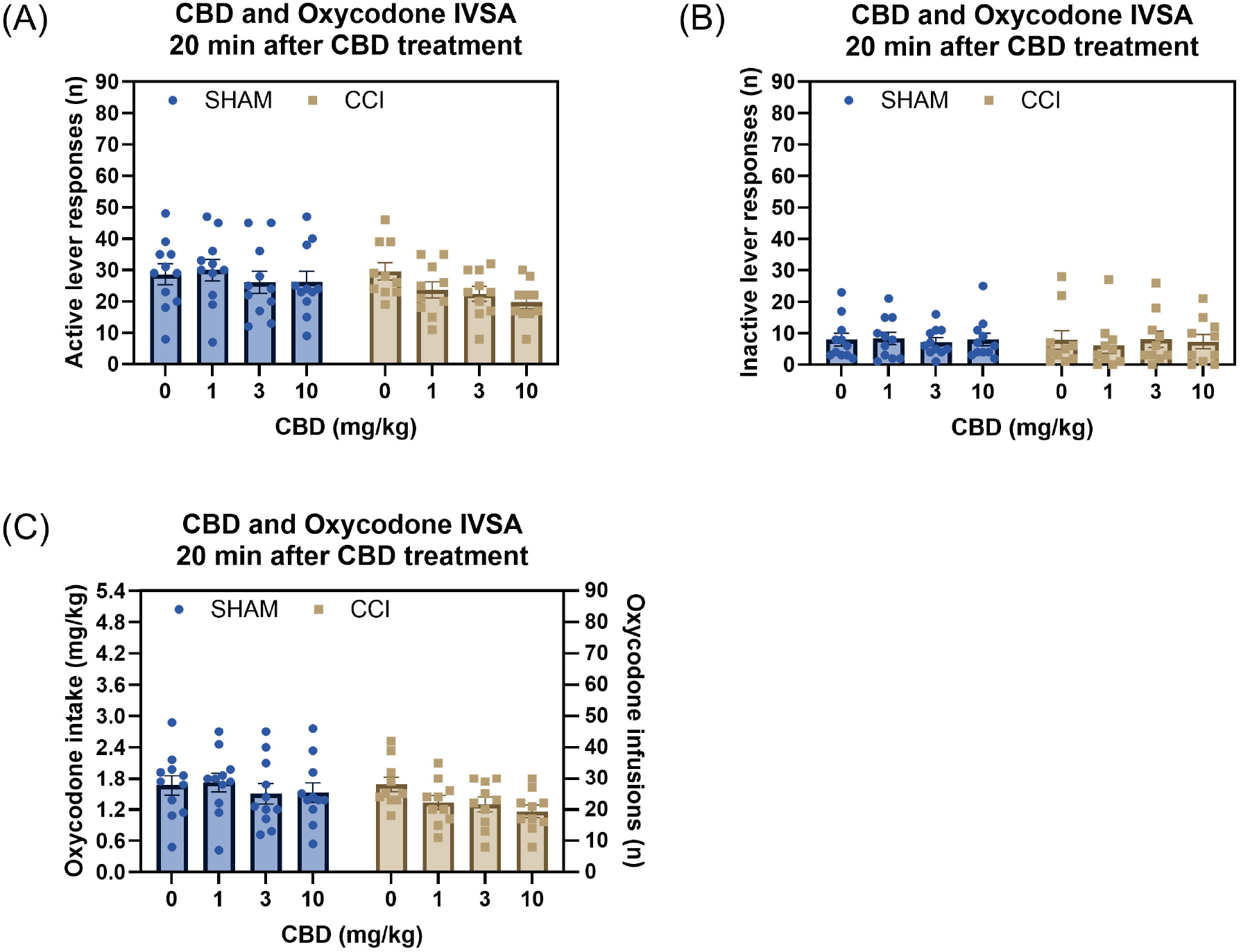
CBD treatment reduces oxycodone intake in both sham control and CCI rats. Treatment with CBD reduced active lever responses in both the CCI and sham control groups, with no effect of CCI surgery on active lever responses or the response to CBD (A). Neither CBD treatment nor CCI surgery affected inactive lever responses (B). CBD treatment decreased oxycodone intake in both the CCI and sham control groups, while CCI surgery had no effect on oxycodone intake (C). Sham, N=11; CCI N=10. Data are presented as mean ± SEM.

#### 3.4.2. CBD and Oxycodone-Induced Antinociception (Hargreaves Test)

##### Absolute paw withdrawal latency

In drug-free condition (baseline), CCI surgery decreased paw withdrawal latency in the surgery paw (Surgery F1,19=5.541, P < 0.05, Paw F1,19=17.54, P < 0.001; Paw × Surgery F1,19=15.388, P < 0.001). However, immediately following oxycodone self-administration, this reduction in latency was reversed in both vehicle and CBD treated CCI rats (Fig. 6A; Treatment F4,76=3.475, P < 0.05; Treatment × Surgery F4,76=1.219, NS; Treatment × Paw F4,76=1.84, NS; Treatment × Paw × Surgery F4,76=2.597, P < 0.05). The post hoc analysis revealed that the significant reduction in withdrawal latency in the surgery paw compared with the control paw was only observed during the baseline drug-free condition in CCI rats. In addition, post hoc analysis showed that oxycodone self-administration had no effect on paw withdrawal latency in either vehicle or CBD treated sham rats.

**Figure 6.**
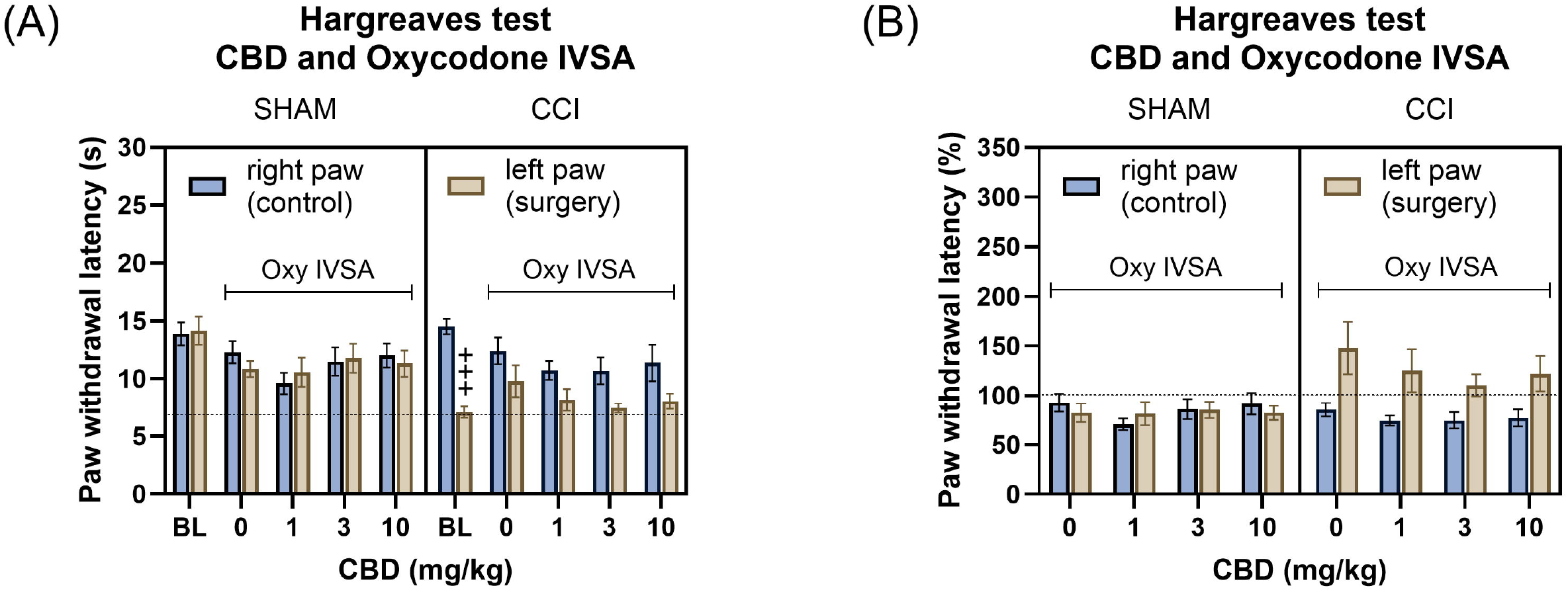
CBD treatment did not alter the antinociceptive effect of oxycodone self-administration in CCI rats. The Hargreaves data are shown as absolute paw withdrawal latency (A) and as percentage change from baseline (B). CCI surgery reduced paw withdrawal latency in the ipsilateral (surgery) paw compared to the contralateral (control) paw, indicating increased thermal sensitivity in the Hargreaves test (A). In vehicle-treated CCI rats, oxycodone self-administration increased paw withdrawal latency in the CCI paw compared to the contralateral paw, indicating antinociceptive effect (A and B). However, treatment with CBD in combination with oxycodone self-administration did not alter the paw withdrawal latency in in the CCI paw compared to the contralateral paw (A and B). Plus signs indicate significantly lower paw withdrawal latencies in CCI surgery paw compared to the contralateral paw in CCI rats. BL, baseline drug-free Hargraves data; Oxy, oxycodone. +++ P<0.001. Sham, N=11; CCI N=10. Data are presented as mean ± SEM.

##### Percentage change in paw withdrawal latency from baseline

To assess the relative changes from baseline, paw withdrawal latencies were expressed as percentage change (Fig. 6B). In CCI rats, oxycodone self-administration increased paw withdrawal latency in surgery paw (Fig. 6B, Surgery F1,19=3.113, NS; Paw F1,19=7.8, P < 0.05; Paw × Surgery F1,19=9.484, P < 0.01). CBD treatment did not alter the effect of oxycodone self-administration on paw withdrawal latency in the CCI surgery paw (Fig. 6B, Treatment F3,57=1.62, NS; Treatment × Surgery F3,57=1.007, NS; Treatment × Paw F3,57=0.601, NS; Treatment × Paw × Surgery F3,57=0.962, NS).

### 3.5. CBD and oxycodone self-administration: 24 hours after treatment

During the self-administration sessions interspersed between CBD test sessions, neither treatment with CBD nor CCI condition affected responding on the active lever (Fig. S1A, Surgery F1,19=0.003, NS; Treatment F3,57=1.44, NS; Treatment × Surgery F3,57=0.408, NS) or the inactive lever (Fig. S1B, Surgery F1,19=0.517, NS; Treatment F3,57=0.422, NS; Treatment × Surgery F3,57=0.511, NS). There was also no effect of CBD treatment or CCI surgery on oxycodone intake (Fig. S1C, Surgery F1,19=0.117, NS; Treatment F3,57=1.469, NS; Treatment × Surgery F3,57=0.103, NS).

## 4. Discussion

In this study, we investigated the effects of CBD and CCI on oxycodone self-administration and thermal pain sensitivity in adult rats. The rats acquired oxycodone self-administration, as indicated by more lever presses on the active compared to the inactive lever. The CCI procedure reduced the paw withdrawal latency in the Hargreaves test, confirming the development of thermal hyperalgesia, a hallmark of neuropathic pain. However, CCI alone did not alter oxycodone self-administration. Treatment with CBD significantly reduced oxycodone intake in both sham and CCI rats. Notably, oxycodone self-administration produced antinociceptive effects in CCI rats but not in sham rats. Furthermore, CBD did not significantly alter the antinociceptive effects of oxycodone in CCI rats. These findings suggest that CBD reduces oxycodone self-administration regardless of pain status while preserving its antinociceptive effects in neuropathic pain.

CCI of the sciatic nerve is a well-established model of chronic neuropathic pain in rodents (Baliki et al., 2005; Guillemette et al., 2012; Jørgensen et al., 2012; Wang et al., 2018; Sheehan et al., 2021; Xi et al., 2023). In this study, CCI decreased the withdrawal latency in the Hargreaves test, indicating that CCI induces thermal pain hypersensitivity, a feature of chronic neuropathic pain. Interestingly, we observed no effect of CCI surgery on oxycodone self-administration, which contrasts with some previous reports. For instance, it has been reported that spinal nerve ligation of the L5 and L6 dorsal nerve roots reduces opioid (heroin, morphine, fentanyl, hydromorphone, and methadone) self-administration in rats (Martin et al., 2007). Similarly, rats with spared nerve injury, which involves ligation of the left tibial and common peroneal nerves while sparing the sural nerve, show reduced fentanyl self-administration compared to sham-operated controls (Rivera-Garcia et al., 2023). Furthermore, a study using a chronic inflammatory pain model (intraplanar injection of Complete Freund’s Adjuvant, CFA) found that pain modulated how heroin affected dopamine release in the nucleus accumbens and heroin self-administration, with these effects being dose-dependent (Hipólito et al., 2015). A low dose (0.075 mg/kg, IV) of heroin reduced dopamine release, whereas a high dose (0.15 mg/kg, IV) enhanced dopamine release. Inflammatory pain did not affect the self-administration of 0.05 and 0.1 mg/kg/infusion of heroin but increased the intake of the higher dose (0.2 mg/kg/infusion)(Hipólito et al., 2015). Taken together, these studies suggest that different types of opioids and the doses of opioids used in chronic pain conditions may differently affect the self-administration of opioids. In the current study, we examined the effects of neuropathic pain on oxycodone self-administration using a single dose (0.06 mg/kg/infusion) in CCI rats. However, previous studies have shown that rats readily self-administer higher doses of oxycodone (Kimbrough et al., 2020; Nguyen et al., 2021). Therefore, further investigation is warranted to determine whether higher doses of oxycodone may reveal different effects of neuropathic pain on oxycodone self-administration in rats.

In this study, CBD treatment reduced oxycodone intake in both sham and CCI groups. This finding that CBD reduces oxycodone intake aligns with previous studies exploring the effects of CBD on opioid intake. For example, Nguyen et al. demonstrated that acute CBD vapor inhalation decreases oxycodone self-administration, suggesting that CBD may reduce the rewarding properties of oxycodone (Nguyen et al., 2023). Our results also align closely with those of Rivera-Garcia et al., who demonstrated that high-CBD cannabis vapor (64.2% CBD and 7.1% THC) reduces fentanyl self-administration in both spared nerve injury and sham-operated female rats (Rivera-Garcia et al., 2023). Furthermore, both intraperitoneally administered CBD and inhalation of high-CBD whole-plant cannabis extract prevents the development of morphine-induced conditioned place preference in mice and rats (Markos et al., 2018; Rivera-Garcia et al., 2023). In addition, CBD blocked the reward-enhancing effect of morphine in the intracranial self-stimulation procedures in rats (Katsidoni et al., 2013). Inhaled CBD also prevented fentanyl-induced conditioned place preference in mice (Bhandari et al., 2024). Taken together, these findings suggest that CBD decreases the rewarding properties of opioids and mitigate opioid misuse irrespective of pain status. Although it was not the focus of this study, we speculate that CBD may reduce oxycodone intake through interactions with several receptor systems involved in reward and substance use. CBD acts as a negative allosteric modulator at CB1 receptors, potentially reducing oxycodone-induced dopamine release and its rewarding effects (Laprairie et al., 2015; Iyer et al., 2022). Furthermore, CBD is a serotonin 5-HT1A receptor agonist and could thereby decrease the rewarding properties of oxycodone (Katsidoni et al., 2013). Finally, the effects of CBD on the GABAergic system may contribute to its potential to reduce oxycodone intake (Bakas et al., 2017b).

In this study, oxycodone self-administration in vehicle-treated rats increased the paw withdrawal latency in the CCI paw but not in sham rats, suggesting that oxycodone mediated antinociceptive effects only in neuropathic pain conditions. Notably, both sham and CCI rats self-administered similar amounts of oxycodone per session (∼1.6 mg/kg total intake/session), raising the possibility that the lack of effect in the sham rats may be related to insufficient total drug exposure. A previous study reported that oxycodone self-administration at a comparable unit dose (0.06 mg/kg/infusion) produced antinociceptive effects in SD rats in the tail-flick assay (Zhang et al., 2020). However, the total oxycodone intake in that study was higher (∼2.7 mg/kg over a 3-hour session) compared to the intake in our sham group (∼1.6 mg/kg over a 2-hour session)(Zhang et al., 2020). This difference in cumulative intake may be critical, as the median effective dose (ED50) for antinociception following subcutaneous administration of oxycodone in male rats is approximately 1.7 mg/kg, with full antinociception observed at 5.6 mg/kg (Austin Zamarripa et al., 2018). Similarly, the ED_50_ following intraperitoneal administration is about 1.46 mg/kg, with full antinociception at 4 mg/kg (Holtman and Wala, 2006). Unlike a single bolus injections, intravenous self-administration results in cumulative drug exposure that depends on session duration and intake behavior. Therefore, it is possible that the total oxycodone intake in sham rats was below the threshold required for measurable antinociceptive effects. In contrast, the same dose was sufficient to alleviate thermal hypersensitivity in CCI rats, suggesting that pain state modulates the efficacy of oxycodone self-administration.

We did not examine the effects of CBD alone on thermal pain sensitivity in drug-naïve (e.g., saline self-administration) sham or CCI rats. However, a previous study reported that sub-chronic intraperitoneal administration of CBD (0.3–30 mg/kg) on post-CCI days 22–24 produced antinociceptive effects in the hot plate assay in male CCI rats (Silva-Cardoso et al., 2021). In the present study, CBD treatment (1–10 mg/kg) did not affect the antinociceptive effects of oxycodone self-administration in CCI rats. In contrast, in an acute pain model, previous research has shown that CBD blocked the antinociceptive effect of oxycodone in the hot plate test in mice (Harris et al., 2022). Furthermore, CBD in combination with morphine produced subadditive effects on morphine-induced antinociception in the hot plate assay (Neelakantan et al., 2015). Notably, in CCI rats, intraperitoneal CBD produced anti-allodynic effects in the Von Frey mechanical sensitivity test only at the highest dose tested (30 mg/kg), but not at lower doses (3 and 10 mg/kg)(Jesus et al., 2022). Moreover, combining the effective (30 mg/kg) and subeffective (10 mg/kg) doses of CBD with subeffective doses of morphine resulted in anti-allodynic effects only when 30 mg/kg CBD was included (Jesus et al., 2022). Taken together, these findings suggest that the interaction between CBD and opioids in modulating pain sensitivity may depend on the specific doses used, and may result in synergistic, additive, or subadditive effects. In the present study, we did not test the 30 mg/kg dose of CBD and used only a single dose of oxycodone for self-administration. Therefore, future studies are warranted to examine the effects of higher CBD doses and varying oxycodone doses in neuropathic pain models.

In this study, the effects of neuropathic pain, CBD, and oxycodone self-administration were examined only in male rats. Previous studies have shown that CCI of the sciatic nerve induces thermal hypersensitivity in both male and female rodents, with no significant sex differences observed (Tall et al., 2001; Sheehan et al., 2021). However, in an acute pain model, intraperitoneal administration of CBD in naïve male and female rats revealed sex differences in antinociceptive effects as measured by the tail-flick assay. Specifically, only the highest dose (30 mg/kg) was effective in males, whereas females exhibited dose-dependent (0.3, 3, and 30 mg/kg) antinociceptive effects (Arantes et al., 2024). In females, these effects were also dependent on the estrous cycle, with significant antinociception observed during late diestrus but not during proestrus (Arantes et al., 2024). Additionally, in acute pain models, the antinociceptive effect of intraperitoneally administered oxycodone has been reported to be greater in females than in males in the tail-flick assay (Holtman and Wala, 2006). Moreover, previous studies have demonstrated sex differences in intravenous oxycodone self-administration in rats (Mavrikaki et al., 2017; Kimbrough et al., 2020). Therefore, further studies are warranted to investigate the effect of sex on CBD and oxycodone self-administration in neuropathic pain models.

In conclusion, our findings demonstrate that CBD reduces oxycodone self-administration in adult rats, regardless of neuropathic pain status, and does not interfere with the antinociceptive effects of oxycodone in neuropathic pain. These results underscore the therapeutic potential of CBD for reducing oxycodone misuse while preserving its antinociceptive effects.

## CRediT authorship contribution statement

**Adriaan W. Bruijnzeel:** Conceptualization, Supervision, Formal analysis, Writing – Original Draft, Visualization, Project administration, Funding acquisition.

**Azin Behnood-Rod:** Investigation, Project administration.

**Ranjithkumar Chellian:** Formal analysis, Investigation, Writing – Original Draft, Visualization.

**Wendi Malphurs:** Investigation, Project administration.

**Robert M. Caudle:** Conceptualization, Writing – Review & Editing.

**Marcelo Febo:** Conceptualization, Writing – Review & Editing.

**Barry Setlow:** Conceptualization, Resources, Writing – Review & Editing.

**Niall P. Murphy:** Conceptualization, Supervision, Project administration, Writing – Review & Editing.

**John K. Neubert:** Conceptualization, Supervision, Funding acquisition, Writing – Review & Editing.

