## Supplemental Figure S1 for "Cannabidiol Reduces Oxycodone Self-Administration While Preserving Its Analgesic Efficacy in a Rat Model of Neuropathic Pain"

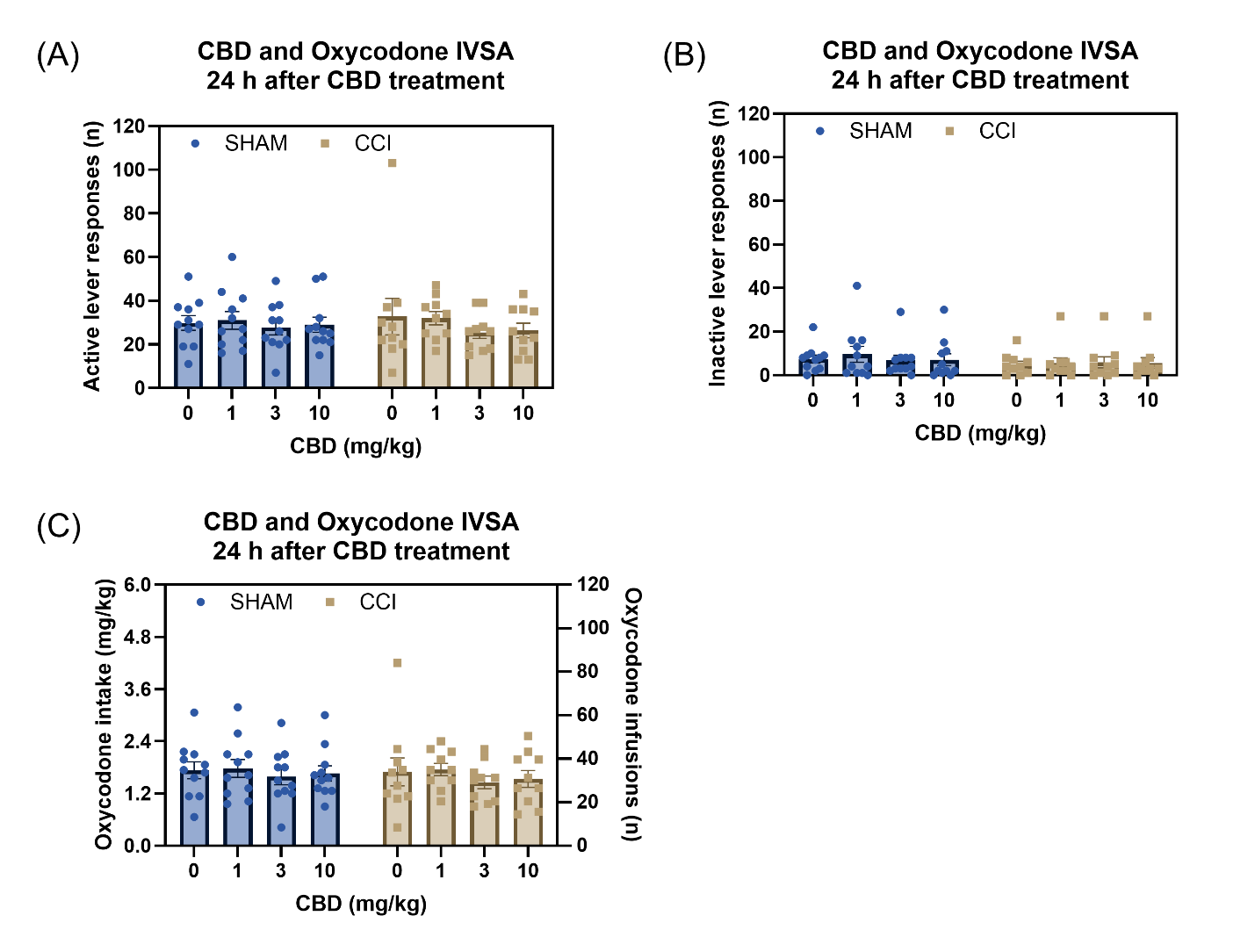

**Figure S1. Effects of CBD treatment and CCI surgery on lever responses and oxycodone intake 24 hours after treatment.** Active lever responses were not significantly affected by CBD treatment or CCI surgery (A). Inactive lever responses were not affected by CBD treatment or CCI surgery (B). Oxycodone intake 24 hours after treatment with CBD was not significantly different between CBD treatment groups or between sham control and CCI surgery groups (C). Sham, N=11; CCI N=10. Data are presented as mean ± SEM.
